# Development of a simple and highly sensitive method for the concentration and detection of African swine fever virus in oral fluids and its feasibility assessment on raised pigs in Northern Vietnam

**DOI:** 10.1101/2024.01.13.575502

**Authors:** Ngan Mai Thi, Giang Tran Thi Huong, Hieu Dong Van, Phan Le Van, Le Huynh Thi My, Dao Bui Tran Anh, Ryoko Uemura, Yasuko Yamazaki, Lan Nguyen Thi, Wataru Yamazaki

**Author notes:** **Address for correspondence:** Wataru Yamazaki, Center for Southeast Asian Studies, Kyoto University, 46 Shimoadachi-cho, Yoshida, Sakyo-ku, Kyoto 606-8501, Japan. These authors contributed equally to this study (Ngan Mai Thi, Giang Tran Thi Huong).

## Abstract

**Background:** Early detection and early slaughter through quarantine are essential to prevent the spread of the African swine fever virus (ASFV). Highly accurate testing is effective for early detection, but it is still difficult to establish a system, especially in the Global South. Pooled oral fluids tests have been used for simple pathogen monitoring, but compared to blood tests, the virus concentration in oral fluids is low, resulting in false negative and missing true positive cases. We collected oral fluids from raised pigs in northern Vietnam and attempted a highly sensitive ASFV survey using a newly developed virus concentration and detection method.

**Results:** In a spike test, the developed method showed up to 100 times greater sensitivity than a reference method. To compare and evaluate the performance of the developed method, a total of 68 pooled oral fluid samples were collected, 63 from northern Vietnam and 5 from southern Japan. Using real-time PCR, 9/68 (13.2%) were positive by the reference method, and 23/68 (33.8%) by the developed method. Using real-time LAMP, 1/68 (1.5%) were positive by the reference method and 6/68 (8.8%) by the developed method.

**Conclusions:** The developed method improved the sensitivity of ASFV detection from oral fluids and enabled early diagnosis of pigs before the onset of the disease. The developed method has the potential to enable simple and highly sensitive diagnosis of ASF, which is essential for its early diagnosis and effective surveillance.

**Highlight:**

- A method has been developed to detect ASFV in pig oral fluids with up to 100 times greater sensitivity than a reference method.
- The developed method combines innovative pretreatment in less than 60 min with conventional nucleic acid extraction with a column kit followed by real-time PCR or LAMP.
- Applied to feasibility assessment for ASF diagnosis on raised pigs, the developed method showed higher diagnostic sensitivity than the reference method.
- The developed method can be a useful tool that contributes to early detection and surveillance of ASFV.

## Introduction

Globalization is accelerating the risk of the worldwide spread of animal infectious diseases (transboundary animal diseases, TADs) [WOAH, World Organisation for Animal Health (formerly known as OIE), 2020, 2021]. Furthermore, TADs are causing serious global economic damage to the livestock and meat industries (WOAH, 2020, 2021) It is therefore important to minimize the spread of pathogens through accurate and highly sensitive diagnosis of TADs (Beemer *et al*., 2019; Howson *et al*., 2018; Yamazaki *et al*.; 2019). One of the bottlenecks, however, is the lack of sophisticated methods to detect pathogens in a simple and sensitive manner.

African swine fever (ASF) is currently a pandemic in Africa, Eurasia, the Dominican Republic and Haiti in the Caribbean, and Papua New Guinea in Oceania, and there is concerned to invade other clean regions such as Japan, the Americas and the Australian continent. ASF is a hemorrhagic disease of domestic and wild suids caused by ASFV infection that is characterized by high fever, hemorrhages in the reticuloendothelial system, and high mortality (Dixon *et al*., 2020; Galindo and Alonso, 2017; Sánchez-Vizcaíno *et al*., 2015; WOAH, 2020, 2021).

Although ASF outbreaks had previously been limited in sub-Saharan Africa, an outbreak in Georgia in 2007 and its subsequent spread to neighboring countries in the south Caucasus, Europe, and the Russian Federation has highlighted the threat of ASF’s transboundary spread (Rowlands *et al*., 2008; Sánchez-Vizcaíno *et al*., 2015; Sauter-Louis *et al*., 2021). Furthermore, ASF invaded China, the world’s largest pig producer in 2018, and then Southeast Asian countries, including Vietnam, continuing to have a devastating impact on the livestock industry and food security (Blome *et al*., 2020; Le *et al*., 2019; Wang *et al*., 2020; WOAH, 2020; You *et al*., 2021; Zhou *et al*., 2018; Zhu *et al*., 2022). As a result of the expansion of ASF, the pig industry has been suffering significant economic losses in ASF-endemic countries (Carriquiry *et al*., 2020; Mason-D’Croz *et al*., 2020; Nguyen-Thi *et al*., 2021; WOAH, 2020; You *et al*., 2021).

After infection with ASFV, pigs are known to shed this virus not only in their blood and organs but also in oral fluids, nasal secretions, feces, and urine (Davies *et al*., 2017; Walczak *et al*., 2020). Among them, oral fluids containing oral fluids has been widely used for monitoring tests such as foot-and-mouth disease virus (FMDV), porcine circovirus type 2 (PCV2), porcine reproductive and respiratory syndrome virus (PRRSV), swine influenza virus (SIV) and vesicular stomatitis virus (VSV) because of its easiness of collection and cost-effectiveness for testing (Beemer *et al*., 2019; Pricket *et al*., 2010; Ramirez *et al*., 2012). However, this method has the disadvantage of causing false-negative results for true positive samples due to insufficient detection sensitivity, since the virus concentration in oral fluids is generally lower than that in blood (Davies *et al*., 2017; Pricket *et al*., 2010; Walczak *et al*., 2020).

The immunomagnetic bead method using specific antibodies has been widely used for enrichment of trace viruses (Dhumpa *et al*., 2011; Makino *et al*., 2020; Yamazaki *et al*., 2019). Although this method is very sensitive, it has the disadvantages of requiring the production of antibodies specific for each virus species and of decreasing LOD due to inhibitory substances derived from the sample. To overcome this problem, we have been already developed a new method for concentration and detection of SARS-CoV-2 in pooled human saliva with 100 times higher sensitivity than a reference method (Yamazaki *et al*., in submission). This developed method, which takes less than one hour to complete, is based on the simple principle of oral fluids lysis by semi-alkaline protease (SAP), a sputum dissolving agent, and concentration of virions by polyethylene glycol (PEG) and high-speed centrifugation, followed by extraction of nucleic acids by a commercial column kit. In this study, we applied this method for the first time to the detection of trace viruses in animal oral fluids and evaluated the feasibility of this method in the sensitive detection of ASFV using 68 pooled oral fluids collected from raised pigs in Vietnam and Japan.

## Materials and Methods

### Oral fluids sampling from raised pigs

A total of 68 samples of pooled oral fluids from raised pigs were tested. Of these, 63 samples obtained from five farmers in four provinces in northern Vietnam between November 20, 2022 and August 25, 2023 were used to validate positive samples. In addition, five samples obtained from five farmers in Miyazaki Prefecture in southern Japan on November 12, 2023 were used as negative controls (Table 1). Pig oral fluids were collected with a cotton rope hung for 30 min in a piggery according to the method described by Ramirez *et al*. (2012), and then, transported to the laboratory. All collected oral fluid samples were stored at −80°C in Vietnam and −30°C in Japan until required. At the time of testing, collected oral fluids were thawed at room temperature, centrifuged at 2,000 *g* for 1-5 min to remove residues of cotton rope and feed, and the resulting oral fluids supernatant was transferred into a new 50 ml sterile plastic tube for both reference and developed methods below described. At this time, no more than 75% of the total oral fluid volume was collected as supernatant, taking care not to inhale any foreign matter at the bottom of the tube. For the spike test to determine the limit of detection (LOD), pig oral fluids were collected from four piggeries in Vietnamese farmers who were confirmed ASF-negative. After mixing 20-30 ml of each pig oral fluids, at least 100 ml of pooled pig oral fluids was used for the spike test as described below.

**Table 1.**
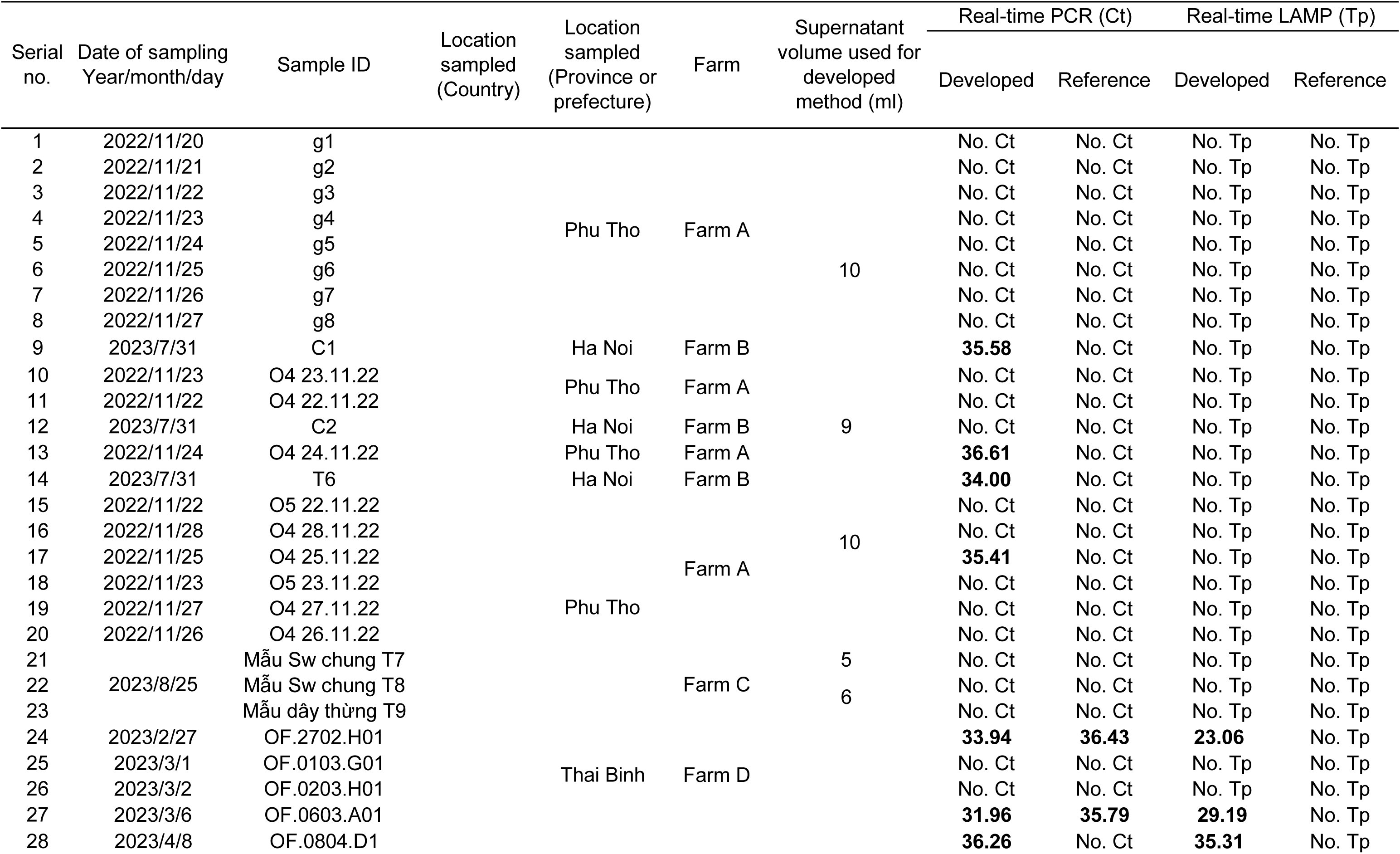
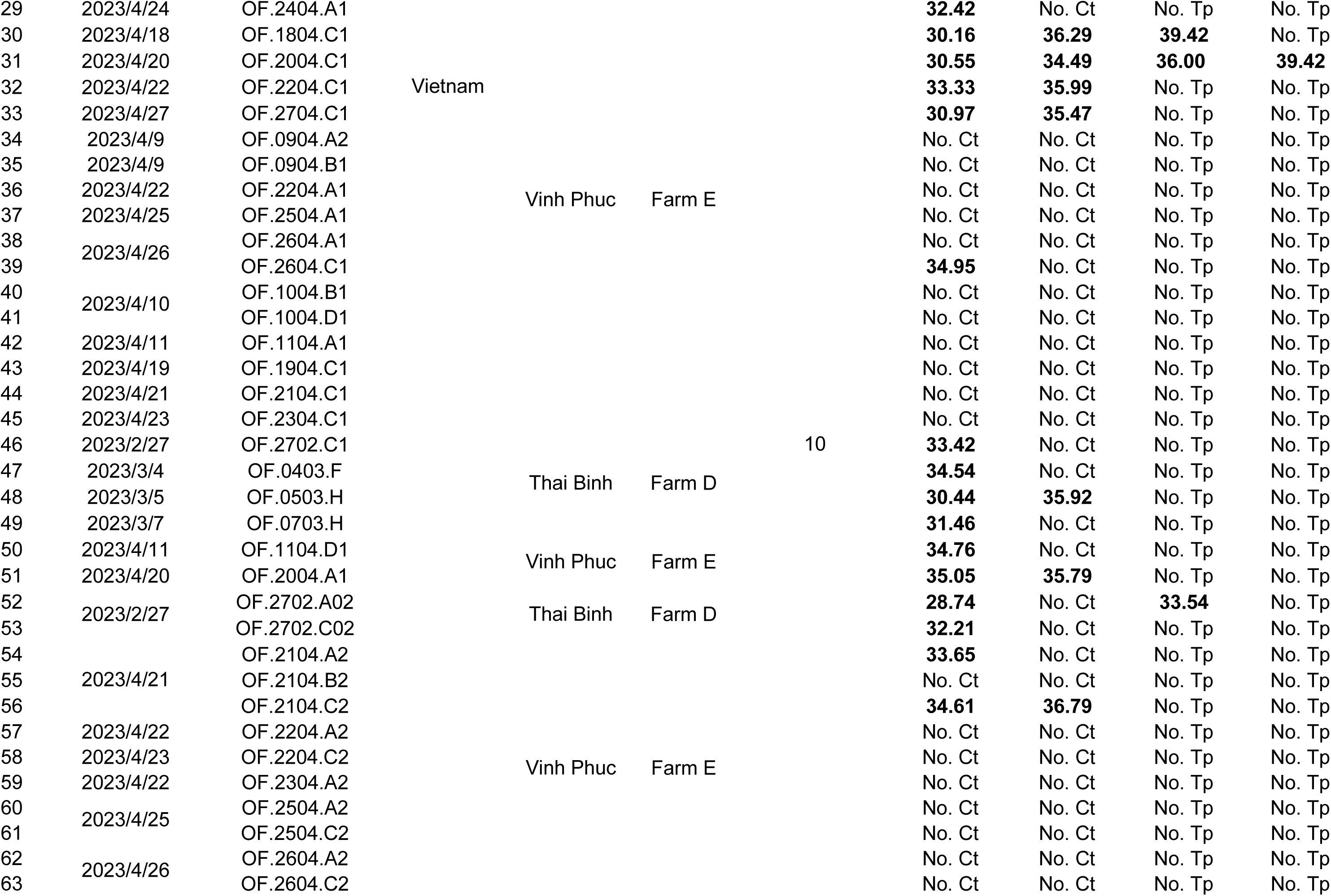
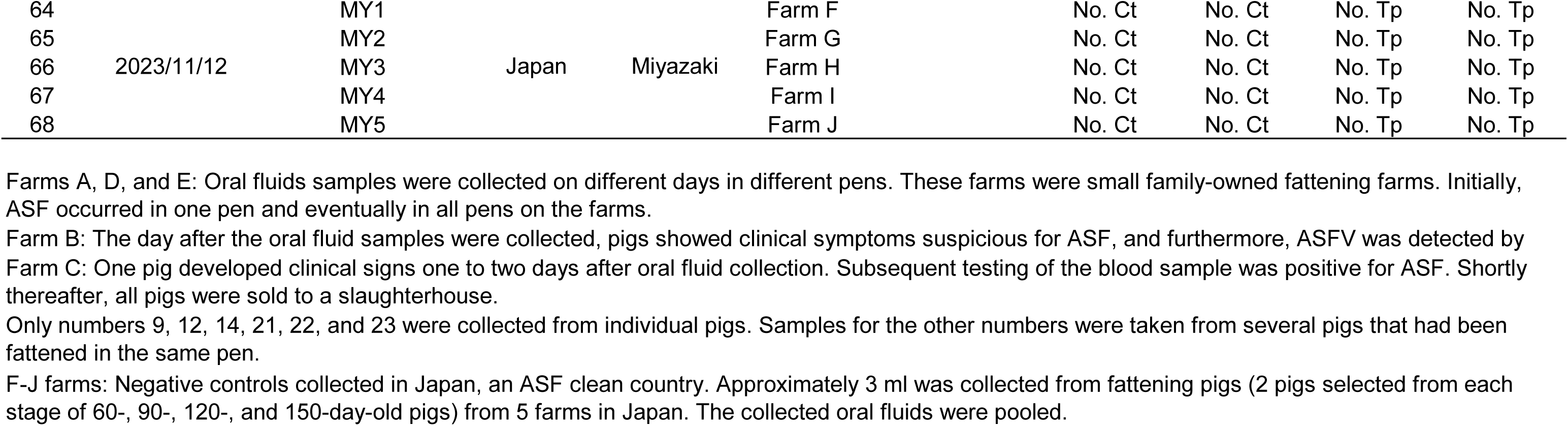
Details of the samples and real-time PCR and LAMP results.

### DNA extraction by reference method

To reduce the viscosity of the oral fluids and prevent clogging of the DNA extraction column in accordance with the pathogen detection manual 2019-nCoV issued by the National Institute of Infectious Diseases, Japan (NIID-J, 2020), 150 μl of pig oral fluids was diluted 1:3 with 450 μl of phosphate-buffered saline (PBS) in a 1.5 ml-sterile microcentrifuge tube. After vertexing, the diluted oral fluids samples were centrifuged at 15,600 *g* for 30 min, and the supernatant was transferred to a new 1.5 ml-sterile microcentrifuge tube. Then, 150 μl of the supernatant was extracted with a DNeasy blood & tissue kit (Qiagen, Maryland, USA), and were extracted as purified DNA in 50 μl of distilled water, otherwise, 200 μl of supernatant was eluted in 50 μl of purified water using the mag LEAD 6gC automated extraction platform (Precision System Science Co., Matsudo, Japan) and a reagent cartridge (MagDEA Dx SV, Precision System Science).

### ASFV concentration by developed method

The method we newly developed to concentrate SARS-CoV-2 from human saliva (Yamazaki *et al*. in submission) was applied to concentrate ASFV from pig oral fluids with slight modification. In brief, 10 ml of pig oral fluid supernatant was placed in a 50 ml-sterile plastic centrifuge tube and diluted 1:3 with 20 ml of SAP (Semi-alkaline proteinase, Suputazyme; Kyokuto Pharmaceutical Industries, Tokyo, Japan). After vortexing, the tubes were kept at room temperature for 15 min to digest oral fluid components. Then, they were centrifuged at 15,557 *g* for 30 min, The supernatant of 24 ml, equivalent to 80% of the total liquid volume, was transferred to a new 50ml-sterile plastic tube. After adding 9.6ml of PEG-NaCl solution equivalent to 40% of the total liquid volume to the new 50 ml-sterile plastic tube (See details of the components of PEG-NaCl solution in our previous publication, Yamazaki *et al*., 2022). The tubes were then vortexed and quickly centrifuged at 8,000 *g* for 20 min to precipitate the ASF virions. after the second centrifugation, the supernatant was carefully removed, and 100 μl (when using the DNeasy blood & tissue kit) or 150 μl (when using the automated extraction platform) was added to the tube with a sterile tip and combined with approximately 50 μl of liquid remaining in the tube to bring the total volume to approximately 150 μl or 200 μl. To detach the ASF virion-PEG complex precipitates, 10 times pipetting were performed, targeting the location on the inner wall of the tube where the ASF virion-PEG complex precipitates were expected to be attached (i.e., where the pellet consisting of oral fluid components was attached at the first centrifugation). A 1 ml-sterile short tip was also added and vortexed to completely detach the pellet. After collecting the mixture in the tube containing the ASF virion-PEG complex by flushing, it was extracted and purified as 50 µl of DNA using an extraction column kit or an automated extraction platform as described above. For the four samples (nos. 12, 21-23, shown in Table 1) whose oral fluid volume was less than 10 ml, the amounts of SAP and PEG-NaCl solution to be added were reduced according to the mixing ratio and tested.

### Real-time PCR assay

Real-time PCR was performed on a CFX Opus 96 Real-Time PCR System (Bio-Rad Laboratories, Hercules, CA, U.S.A.) or a QuantStudio 3 (Thermo Fisher Scientific, Inc., Waltham, MA, U.S.A.) with two primers and probes consisting of the forward primer (5’-TGC TCA TGG TAT CAA TCT TAT CG −3’), the reverse primer (5’-CCA CTG GGT TGG TAT TCC TC −3’) and the probe (5’-FAM-TTC CAT CAA AGT TCT GCA GCT CTT-TAMRA-3’) reported by Tignon and colleagues (2011). Amplification cycle conditions, reagent volumes and concentrations were slightly modified as follows. Briefly, 50 µl of real-time PCR reaction mixtures were used, comprising 25 µl of 2x probe qPCR mix (Takara Bio Inc., Otsu, Japan), 0.2 µl each of forward and backward primers (100 µmol l^-1^ each; Hokkaido System Science, Co., Ltd., Sapporo, Japan), 0.1 µl of probe (100 µmol l^-1^; Hokkaido System Science), 19.5 µl of nuclease-free water (Takara Bio), and 5 µl of the DNA template. The cycling conditions were as follows: one cycle at 95°C for 30 sec, followed by 45 cycles each at 95°C for 10 sec, and 60°C for 60 sec. The automatically calculated Ct value was adopted, and the Ct cut-off value was set at 37.00.

### Real-time LAMP assay

Real-time LAMP assay was performed following a previous report (Mai, *et al*., 2023) with slight modifications. Briefly, in-house LAMP reaction mixtures were prepared by doubling the volume of reaction solution to 50 µl with the same composition as described. The amount of template DNA was set to 5 µl. The reaction mixtures were incubated at 63°C for 45 min, followed by 80°C for 5 min to complete the reaction using a real-time turbidimeter (Loopamp EXIA; Teramecs, Kyoto, Japan). When the derivative of turbidity reached 0.05 within 40 min, the reaction was automatically considered positive. Endpoint judgement by unaided eye was also performed. When the reagent color remained purple, the result determined as negative. On the other hand, a change to sky blue was interpreted as positive. For both real-time PCR and LAMP analysis, positive controls were DNA sequences derived from field isolates in Vietnam and artificial synthetic DNA sequences ordered from Eurofins Genomics K. K. (Tokyo, Japan) in Japan.

### Preparation of 10-fold dilution series of ASFV spiked oral fluids

The ASFV strain ASF/HY01/2019 (GenBank Accession no. MK554698), belonging to p72 genotype II and isolated from infected pigs during the first ASF that occurred in Vietnam in February 2019, was used for the spike test. This strain, which was cultured on porcine alveolar macrophages (PAM) with a TCID_50_ of 106 .5, and stored at −80°C, was thawed at 4°C and used promptly for the spike test. A 10-fold dilution series of the ASFV strain in PBS was prepared and kept at 4°C. In parallel, pooled pig oral fluids stored at −80°C was thawed at room temperature. The supernatant, created by centrifugation at 2,000 *g* for 5 min to remove cotton rope and feed residue, was transferred to two new 50 m-sterile tubes for the spike test. Next, the pig oral fluid supernatant was dispensed into six 50 ml tubes of 10 ml each, and then, the 10-fold dilution series of ASFV was sequentially spiked and vortexed thoroughly. A series of pig oral fluids containing from neat to 10(−5)-fold ASFV dilutions was prepared, as shown in Table 2.

**Table 2.** Limit of detections (LODs) of the developed and reference methods for detecting ASFV spiked into ASFV-negative pig oral fluids.

| Assays | Dilutions of pooled ASFV-spiked oral fluids samples |  |  |  |  |  |
| --- | --- | --- | --- | --- | --- | --- |
|  | 10<sup>-0</sup> | 10<sup>-1</sup> | 10<sup>-2</sup> | 10<sup>-3</sup> | 10<sup>-4</sup> | 10<sup>-5</sup> |
| Real-time PCR (Ct) |  |  |  |  |  |  |
| Developed method | 26.40 | 30.34 | 33.25 | 36.43 | No. Ct | No. Ct |
| Reference method | 35.51 | No. Ct | No. Ct | No. Ct | No. Ct | No Ct |
| Real-time LAMP (Tp) |  |  |  |  |  |  |
| Developed method | 28:36 | 32:38 | No. Tp | No. Tp | No. Tp | No. Tp |
| Reference method | No. Tp | No. Tp | No. Tp | ND | ND | ND |

Ct, Threshold cycle.
Tp, Time of positivity.
No. Ct, No threshold cycle values detected using real-time PCR.
No. Tp, Time-of-positivity values undetectable using LAMP.
ND, Not determined.
\*, In a duplicate analysis, using real-time PCR or LAMP assay, all two samples were positive or negative, with no samples showing one positive and one negative.

### Determination of LODs

The LODs were determined using the pooled pig oral fluids and the 10-fold dilution series of ASFV as described above. DNA was extracted after performing the reference and developed methods described above. In the duplicate analyses by using real-time PCR and LAMP assays, if two samples were positive, the result was interpreted as positive, or if one sample was positive and one sample was negative, or if two samples were negative, the results were interpreted as negative (Table 2).

### Theoretical virion concentration ratio and percentage

The theoretical virion concentration ratio of the developed method was calculated by subtracting the Ct value of the developed method from the Ct value of the reference method, and then, taking the logarithm of 2. That is, the following equation was used: 2^(Ct^ ^reference^ ^method-Ct^ ^developed^ ^method)^. Furthermore, the theoretical virion concentration ratio value was divided by 200, the ratio of the volume of oral fluid supernatant used in the two methods (10 ml in the developed method *vs.* 50 µl in the reference method), and then added to 1 and expressed as a percentage to calculate the theoretical concentration percentage.

## Results

A total of 68 pooled oral fluids were collected from 63 Vietnamese and five Japanese raised pigs, and the performance of the developed method was compared with that of the reference method. As a result, 9/68 (13.2%) of the reference method and 23/68 (33.8%) of the developed method were positive by the real-time PCR assay. By the LAMP assay, the samples showed positive in 1/68 (1.5%) of the reference method and in 6/68 (8.8%) of the developed method (Tables 1, 3, Figures 1 and 2). Among the samples, total of five pooled oral fluids collected from raised pigs in Japan, which were used as negative controls, were all negative, as expected (Table 1). As shown in Table 2, the spike test resulted that the newly developed method was 1000-fold and at least 100-fold more sensitive than the reference method for real-time PCR and real-time LAMP detections, respectively, when pretreatment was applied within 60 min to a 10 ml oral fluid supernatant. The results of the real-time measurement in the LAMP assay and the color change by the unaided eye observation were in perfect agreement (Data not shown).

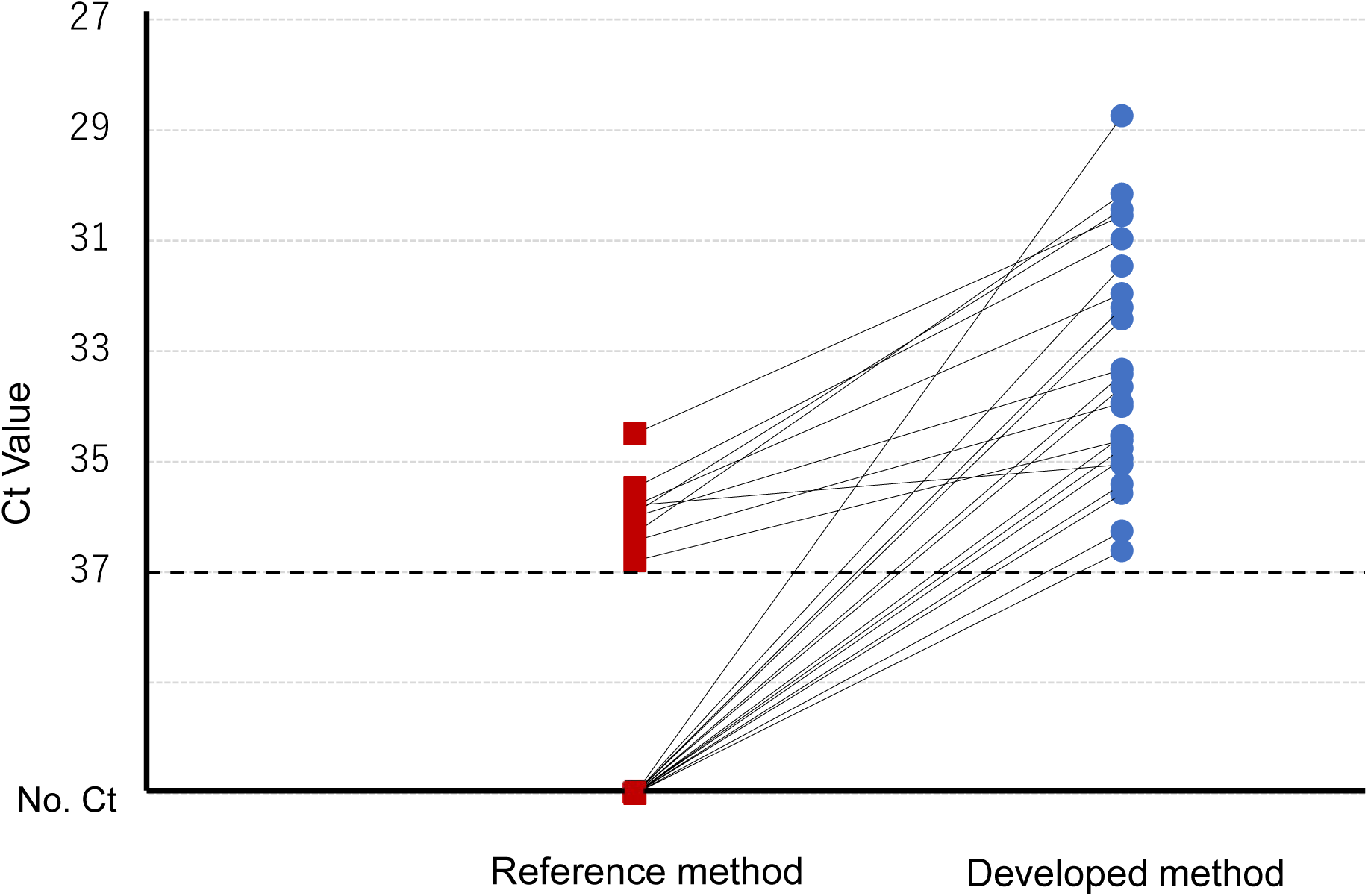

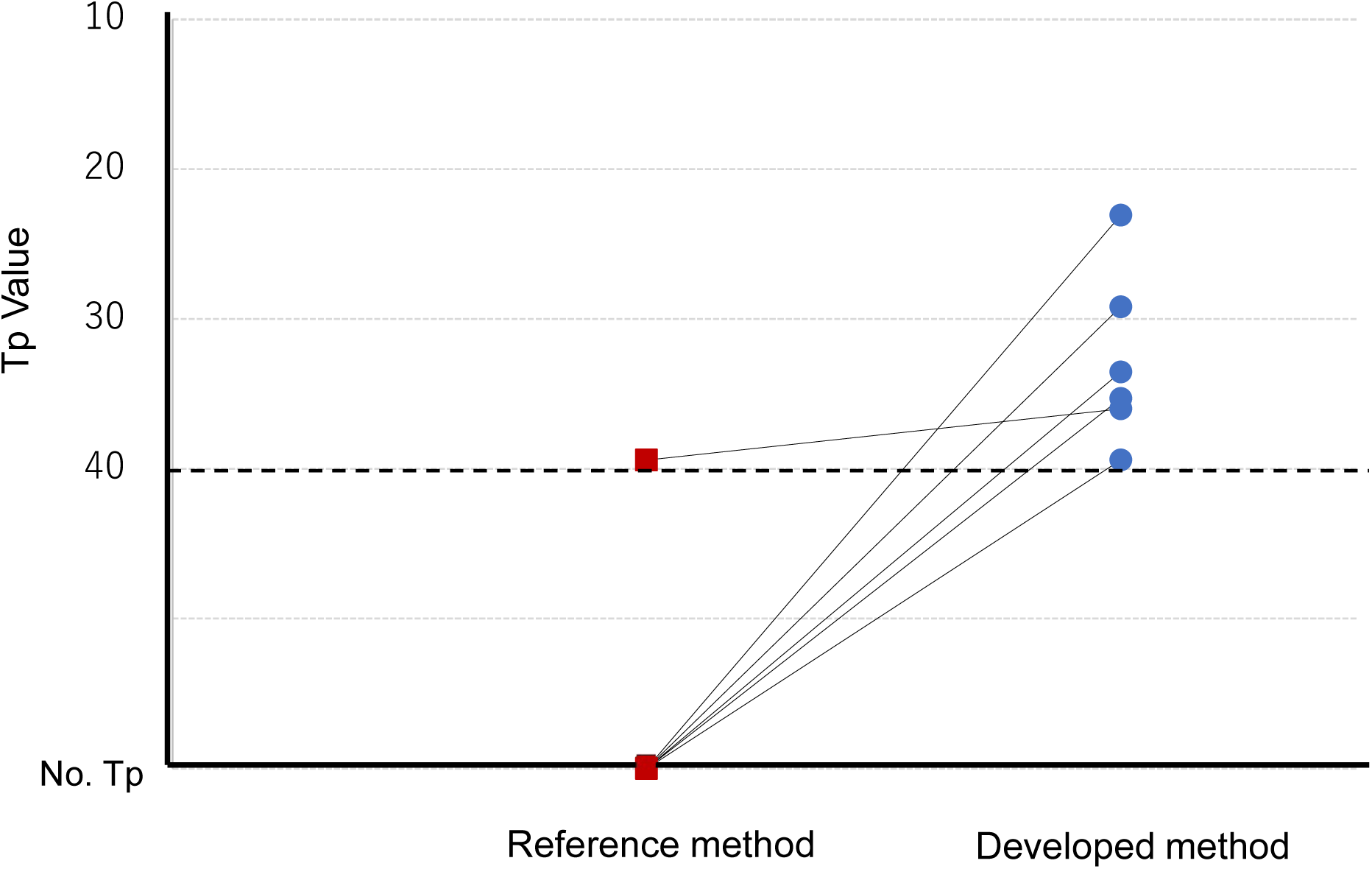

**Table 3.** Diagnostic performance of the developed and reference methods for ASF diagnosis with pig oral fluids using real-time PCR and LAMP assays.

|  | Real-time PCR | Real-time LAMP |
| --- | --- | --- |
| Developed method | 23/63<br>(Sen, 36.5%) | 6/63<br>(Sen, 9.5%) |
| Reference method | 9/63<br>(Sen, 14.3%) | 1/63<br>(Sen, 1.6%) |

Sen, Diagnostic sensitivity.

Based on the positive values obtained and the sample volumes used in both developed and reference methods, theoretical concentration ratio and percentage for the 23 positive samples were calculated and showed values ranging from >1.3 to >306.6 times and >100.7% to >253.3%, respectively (Table 4). However, accurate calculations were difficult because 14 samples were positive only by the developed method, while all 14 were negative by the reference method (Table 4 and Figure 1).

**Table 4.**
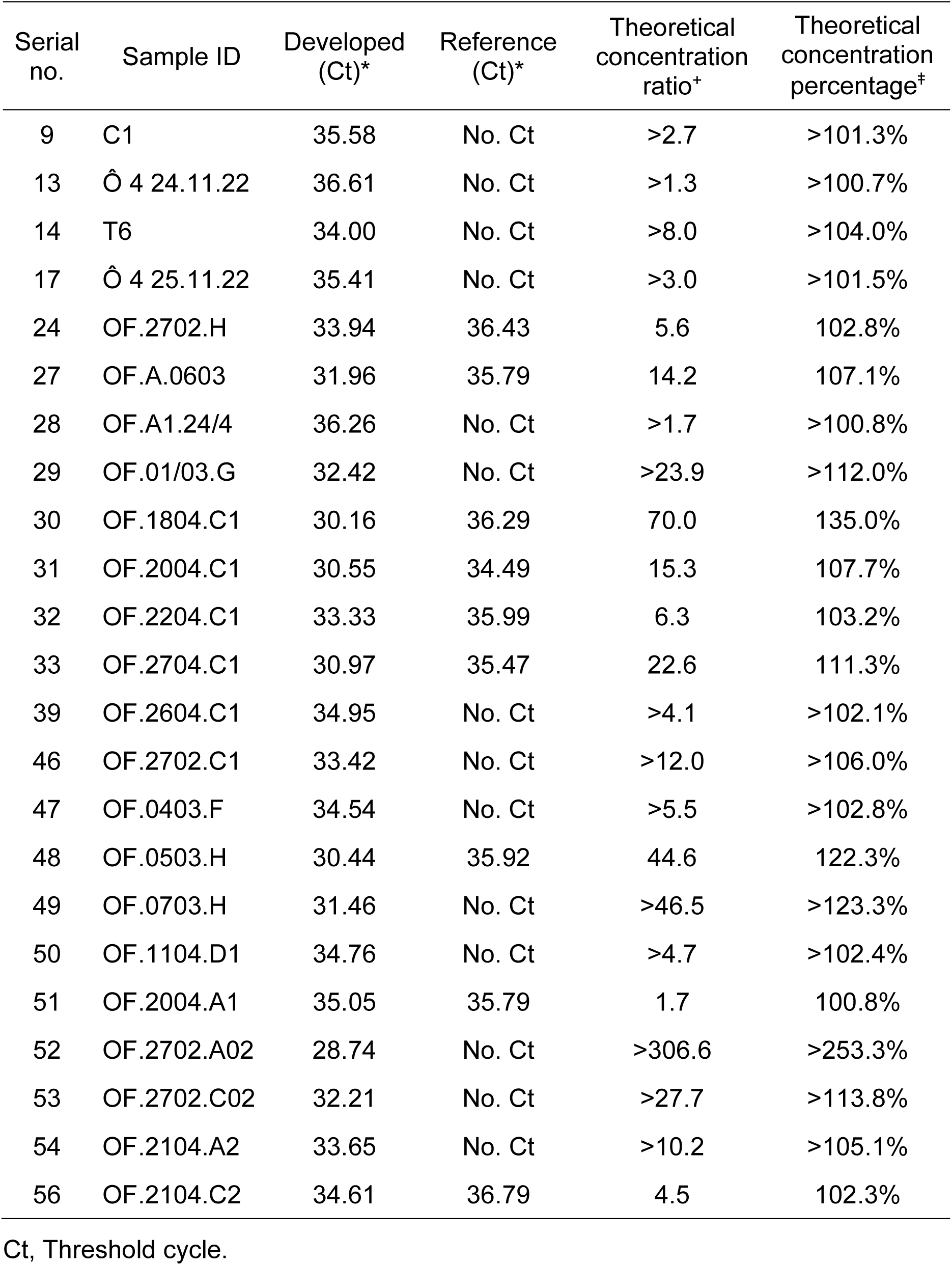
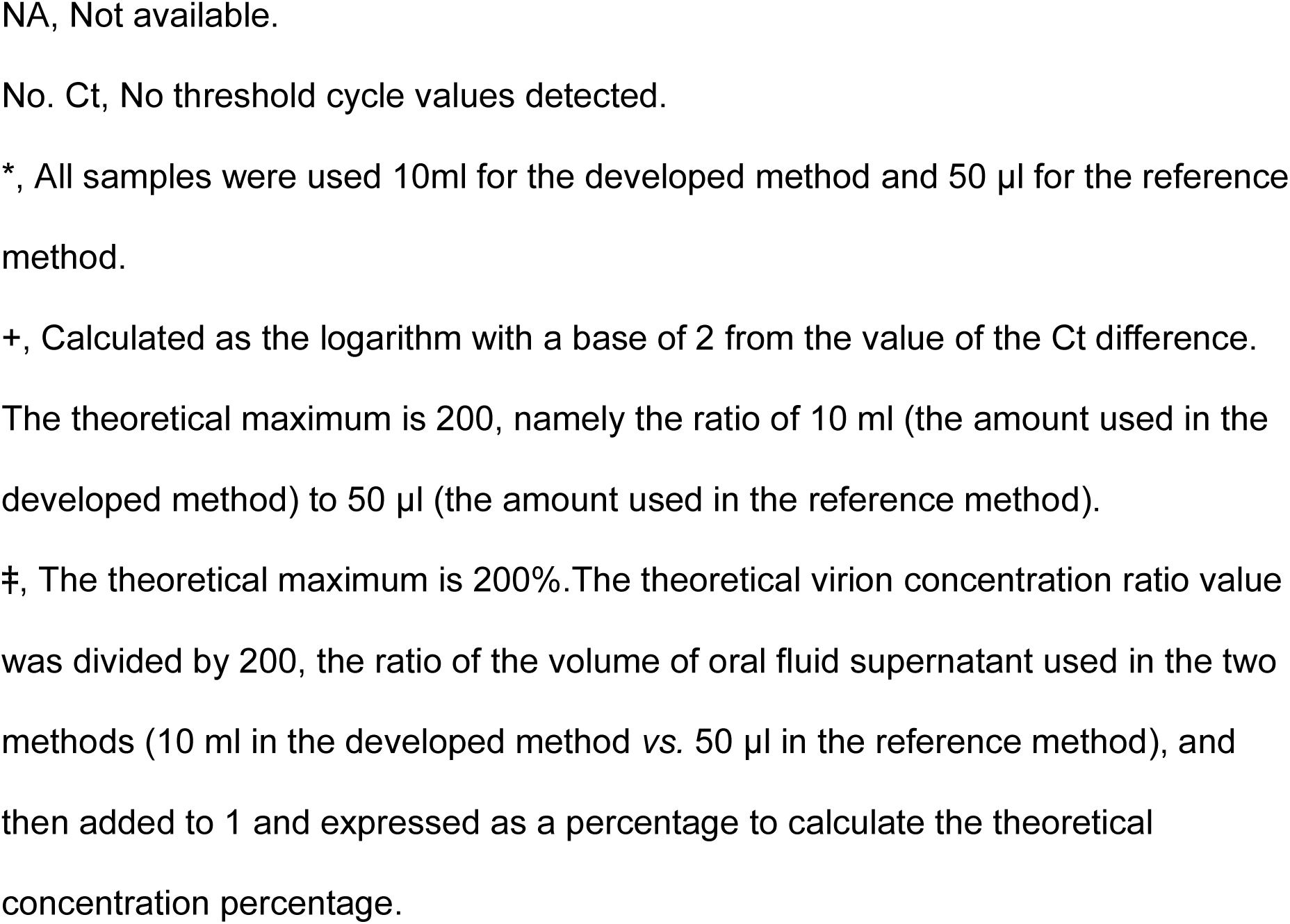
Theoretical concentration and recovery rates of 23 samples that showed positive by real-time PCR detection using the developed or reference method.

| Serial no. | Sample ID | Developed (Ct)* | Reference (Ct)* | Theoretical concentration ratio<sup>+</sup> | Theoretical concentration percentage<sup>‡</sup> |
| --- | --- | --- | --- | --- | --- |
| 9 | C1 | 35.58 | No. Ct | >2.7 | >101.3% |
| 13 | Ô 4 24.11.22 | 36.61 | No. Ct | >1.3 | >100.7% |
| 14 | T6 | 34.00 | No. Ct | >8.0 | >104.0% |
| 17 | Ô 4 25.11.22 | 35.41 | No. Ct | >3.0 | >101.5% |
| 24 | OF.2702.H | 33.94 | 36.43 | 5.6 | 102.8% |
| 27 | OF.A.0603 | 31.96 | 35.79 | 14.2 | 107.1% |
| 28 | OF.A1.24/4 | 36.26 | No. Ct | >1.7 | >100.8% |
| 29 | OF.01/03.G | 32.42 | No. Ct | >23.9 | >112.0% |
| 30 | OF.1804.C1 | 30.16 | 36.29 | 70.0 | 135.0% |
| 31 | OF.2004.C1 | 30.55 | 34.49 | 15.3 | 107.7% |
| 32 | OF.2204.C1 | 33.33 | 35.99 | 6.3 | 103.2% |
| 33 | OF.2704.C1 | 30.97 | 35.47 | 22.6 | 111.3% |
| 39 | OF.2604.C1 | 34.95 | No. Ct | >4.1 | >102.1% |
| 46 | OF.2702.C1 | 33.42 | No. Ct | >12.0 | >106.0% |
| 47 | OF.0403.F | 34.54 | No. Ct | >5.5 | >102.8% |
| 48 | OF.0503.H | 30.44 | 35.92 | 44.6 | 122.3% |
| 49 | OF.0703.H | 31.46 | No. Ct | >46.5 | >123.3% |
| 50 | OF.1104.D1 | 34.76 | No. Ct | >4.7 | >102.4% |
| 51 | OF.2004.A1 | 35.05 | 35.79 | 1.7 | 100.8% |
| 52 | OF.2702.A02 | 28.74 | No. Ct | >306.6 | >253.3% |
| 53 | OF.2702.C02 | 32.21 | No. Ct | >27.7 | >113.8% |
| 54 | OF.2104.A2 | 33.65 | No. Ct | >10.2 | >105.1% |
| 56 | OF.2104.C2 | 34.61 | 36.79 | 4.5 | 102.3% |

Ct, Threshold cycle.
NA, Not available.
No. Ct, No threshold cycle values detected.
\*, All samples were used 10ml for the developed method and 50 µl for the reference method.
+, Calculated as the logarithm with a base of 2 from the value of the Ct difference. The theoretical maximum is 200, namely the ratio of 10 ml (the amount used in the developed method) to 50 µl (the amount used in the reference method).
‡, The theoretical maximum is 200%.The theoretical virion concentration ratio value was divided by 200, the ratio of the volume of oral fluid supernatant used in the two methods (10 ml in the developed method vs. 50 µl in the reference method), and then added to 1 and expressed as a percentage to calculate the theoretical concentration percentage.

## Discussion

We revealed for the first time that a newly developed virion concentration method can be used to reliably detect ASFV in the oral fluids of ASF-infected pigs prior to the onset of ASF, leading to early detection. This simple developed method uses a large amount of oral fluids (>10 ml), and thus has a detection sensitivity up to 100 times greater than the conventional reference method, which uses only 50-200 μl, making it effective in strengthening the ASF survey and early detection system worldwide. Although >10 ml of oral fluids can easily be collected for oral fluids testing of pigs, only 50-200 µl of oral fluids is used for the actual test. In this study, we attempted to improve diagnostic performance by increasing the theoretical concentration rate up to 200 times by using 50 to 200 times the conventional sample volume.

The developed method has shown that it can greatly improve the technical limitation of conventional genetic testing methods, namely, the problem that samples containing trace amounts of virus have been determined to be false-negative because they are below the LOD. Assuming a Ct value below 35.00 as a definite positive and a Ct value of above 35.00 to below 37.00 as a borderline difficult to determine, with the reference method, only 11.1% (1/9) of the ‘positive’ samples were definite positive and the remaining 88.9% (8/9) were borderline, making the determination difficult. In contrast, with the developed method, 78.3% (18/23) of the ‘positive’ samples have Ct values below 35.00, making it possible to determine a positive result with certainty. In other words, the reference method was difficult to determine because the Ct values of the nine "positive" samples ranged between 34.49 and 36.79, and eight of the nine samples were above Ct 35.00. In contrast, with the developed method, the Ct values of the 23 "positive" samples ranged between 28.74 and 36.61, and 18 samples with Ct values below 35.00 could be clearly determined as ‘positive’ except for five samples that were above Ct 35.00 (Tables 1, 3, 4 and Figure 1).

In the LAMP assay, only one sample was positive at 39 min 42 sec in the reference method and six samples were positive at 23 min 6 sec to 39 min 42 sec in the developed method (Table 1 and Figure 2). In contrast, in the real-time PCR assay, nine and 23 samples were positive for the reference and developed method, respectively. As shown in Table 2, the LAMP assay has already been found to be at least 10- to 100-fold less sensitive than the real-time PCR assay in the spike test, as well as in our previous report (Mai *et al*., 2023). Therefore, the low positivity rate by the LAMP assay is rationale and not surprising.

As shown in Tables 1-4, Figures 1 and 2, the concentration of virion by the developed method resulted in an increase in the positivity of both the real-time PCR and LAMP assays. On the other hand, there was a difference between the actual concentration ratio at 200 times (100% recovery rate) and the theoretical concentration ratio between >1.3 to >306.6 times (>100.7% to >253.3% theoretical concentration percentage), and the concentration effect was not as high as shown in the spike test (Tables 2 and 4). This is not surprising, since both values decrease greatly as the amount of the virion in the sample decreases. For example, the theoretical concentration ration and percentage from the spike test are 552.6 times and 376.3% for the 10^-0^ dilution, >101.1 times and >150.6% for the 10^-1^ dilution, >13.5 times and >106.7% for the10^-2^ dilution, and >1.5 times and >100.7% for the 10^-3^ dilution. This is merely a technical limitation that, as with the reference method, the detection threshold of a typical real-time PCR method is set around Ct 37.00, so the apparent theoretical recovery rate is conservatively calculated based on Ct 37.00 as the baseline. In other words, the developed method is highly sensitive even for samples below the LOD of the reference method.

ASFV was also detected in oral fluid collected prior to confirmation of the ASF outbreak (Table 1). The farm was later confirmed positive for ASFV. In particular, on farm B, where oral fluid samples nos. 9, 12, and 14 were collected from individual pigs on July 31, 2022, a blood test was urgently performed because the day after the oral fluid samples were collected (August 1), the pigs showed clinical signs suspicious for ASF. The next day, August 2, i.e., the second day of the oral fluid collection, the blood test was confirmed positive for ASFV. on August 2, when the ASFV positivity was confirmed, the farmer sold all pigs to the slaughterhouse. As oral fluid samples collected on July 31 were negative by the reference method but positive by the developed method (Tables 1 and 3), the developed method enabled early detection of positive pigs without onset. One limitation of this study is that although the concentration process is simple, care must be taken to avoid contamination at the laboratory due to the large number of steps involved.

Apart from the points mentioned above, it is necessary to consider the variation in the values obtained by the developed method, i.e., the fact that some samples, such as nos. 51 and 56, showed very small differences of the Ct values between the developed and the reference methods. One possible reason of this discrepancy may be caused that the spike test used fresh ASFV immediately after incubation, and the oral fluid samples used in the performance evaluation were collected in the field and stored at −80°C until the experiment. In other words, if ASFV is present in pig oral fluids as DNA rather than virion, the detection sensitivity of the developmed method, which is concentrating virion, may be reduced because only the reference method can detect the DNA. Alternatively, in our preliminary experiment, the theoretical recovery rate of virions was sometimes reduced by more than 10-fold due to the increase in Ct value when pellets derived from oral fluids were not completely removed by the centrifugation process and remained visible by the unaided eye observation in the process immediately preceding DNA extraction, the process of collecting the virion-PEG complex from the tubes (data not shown). Since visible pellets derived from oral fluids components were also observed in some samples in this study, this may be the cause of some reduced concentration efficacy.

In our preliminary study, we attempted to detect ASFV using reference and developed methods in six samples of oral fluids from semi-farmed wild boars (a unique form of boar husbandry in which boars graze in the forest during the day and spend the night in the farmer’s pig pen) traditionally raised in mountain villages in Vietnam. However, all the oral fluids samples were negative (data not shown). This is a natural result considering the extremely low ASFV harboring rate in healthy wild boars (Sauter-Louis *et. al*, 2021). In the future, we plan to collect wild boar oral fluids using bite traps and other methods, and then, conduct a study to determine the ASFV harboring in wild boars using the developed method with high sensitivity. This may allow for highly accurate monitoring of the spread of ASFV through wild boars in the environment.

It is clear that the developed method can reliably detect trace amounts of virus in pig oral fluids naturally infected with ASF by using a larger sample volume than before. As a simple, highly sensitive, and cost-effective oral fluids virus test method, it is expected to be applied to highly sensitive diagnosis of various diseases such as foot-and-mouth disease in ruminants and rabies in dogs, as well as infectious diseases in pigs.

## Conclusions

We have succeeded in developing a simple and highly sensitive method for concentrating ASFV in oral fluids from raised pigs and demonstrated that the developed method has extremely sensitive detection performance than conventional methods. We also succeeded in detecting ASFV in oral fluids of naturally infected pigs with higher sensitivity than conventional methods, demonstrating the high feasibility of the developed method.

## Ethics approval

The authors confirm that the ethical policies of the journal, as noted on the journal’s author guideline page, have been adhered to. Samples used in this study were those submitted to the Vietnam National University of Agriculture, Hanoi, Vietnam and University of Miyazaki, Miyazaki, Japan, for ASFV diagnosis. Ethical approval was not required.

## Declaration of Availability of data and materials

All data obtained in this study is included within the paper. In addition, the data sets in this study are available from the corresponding author upon reasonable request.

## Declaration of Submission

The authors confirm that this manuscript or data has not been previously published and is not being considered for publication elsewhere. The authors further confirm that all authors have contributed to the study and have approved the final version.

## Authors’ contributions

**MTN**: Collecting samples, Data curation, Funding acquisition, Formal analysis, Methodology, Visualization, Writing - original draft. **TTHG**: Data curation, Methodology, Writing - original draft. **DVH**: Data curation, Formal analysis, Methodology, Visualization. **Le Van Phan**: Collecting samples, Methodology. **HTML**: Formal analysis, Funding acquisition. **BTAD**: Funding acquisition. **RU**: Collecting samples. **YY**: Data curation, Funding acquisition, Formal analysis, Methodology, Visualization, Writing - original draft, **NTL**: Funding acquisition, Supervision. **WY**: Conceptualization, Data curation, Funding acquisition, Methodology, Project administration, Resources, Supervision, Validation, Visualization, Writing - original draft.

## Abbreviations

ASF: African swine fever
LOD: Limit of detectiony
PBS: Phosphate-buffered saline
PEG: Polyethylene glycol
SAP: Semi-alkaline proteinase

